# A novel mouse model of voluntary stretching and its application in breast cancer research

**DOI:** 10.1101/2025.01.24.634735

**Authors:** Yi He, Lisbeth Berrueta, Yan Wang, Gary J. Badger, Helene M. Langevin

## Abstract

**Background:** Stretching exercises such as yoga are recommended for cancer survivors to manage symptoms and promote wellbeing in clinical settings. Although other types of exercise (e.g. running) can reduce the growth of tumors in animal models, the role of stretching on tumor growth remains unclear, and the lack of a preclinical self-stretching model has impeded mechanistic studies on health benefits of stretching. We sought to develop a voluntary stretching animal model to address this research gap and apply it to breast cancer research.

**Methods:** Using food, water, and enrichment in the home cage as motivators for stretching, a two-week 24/7 behavior monitoring was conducted in a video-based customizable home-cage behavior tracking system, Noldus PhenoTyper, to promote self-stretching in FVB mice. Subsequently, this model was utilized in a comparative study of voluntary stretching and voluntary running on tumor growth and plasma protein profiles in the MET-1 orthotopic mammary tumor FVB mouse model.

**Results:** The new voluntary stretching model effectively elicited mouse self-stretching in the custom cage setting in the long-term observation and significantly inhibited tumor growth as effectively as voluntary wheel running. Moreover, plasma proteomic analysis demonstrated that voluntary stretch versus voluntary running distinctly impacted systemic protein profiles, possibly linking to different cellular and molecular mechanisms underlying anti-cancer effects and, potentially, exercise-induced benefits in other health conditions.

**Conclusion:** Our work provides the first preclinical voluntary stretching model, which may be well suited to breast cancer research and a valuable research tool to facilitate investigations of stretching health benefits across various research fields.

## 1. Introduction

Aerobic, resistance, and flexibility exercises are the three main types of exercises recommended for cancer survivors. According to the American College of Sports Medicine (ACSM), exercise has been demonstrated to prevent seven common cancers and prolong the survival of patients with three cancers, including breast cancer.^1^ Similarly, research using animal exercise models has also shown anti-tumor effects of exercise, with notable evidence for breast cancer.^2^ To date, several aerobic-exercise-based experimental rodent models, such as voluntary wheel running,^3–7^ forced treadmill,^8–11^ and swimming^12–15^ are the primary tools to explore the mechanisms underlying the positive impact of exercise on breast cancer.

However, it is not negligible that some studies based on these exercise models have offered conflicting findings regarding the protective effects of exercise on breast cancer.^16–22^ For instance, high-intensity treadmill running upregulated the expression of tumor proliferation antigen^20^ and accelerated bone destruction by metastasized breast cancer.^21^ Furthermore, forced treadmill training and spontaneous wheel running have also unexpectedly been found to stimulate mammary tumorigenesis,^16–18, 20, 23^ shorten animal survival time,^18^ and even potentiate pulmonary metastasis.^19, 24^ Apart from tumor-model-specific responsiveness to exercise,^25^ the intensity of exercise could be a contributor to these detrimental effects.^4, 19, 22^ In the clinical setting, the intensity of exercise intervention is also a crucial aspect to evaluate before it is prescribed to cancer patients,^26^ due to safety and tolerance concerns associated with cancer status (e.g. bone metastasis) and treatment-induced syndrome (e.g. osteoporosis, fatigue, and pain).^27, 28^ Therefore, cancer patients and survivors are generally advised to avoid excessive high-intensity exercise.^28^

Besides aerobic and strength workouts, low intensity flexibility exercises are one of the three modalities of exercise recommended in the cancer patient exercise guideline.^28–30^ Thus far, the clinical studies of flexibility training, either as a part of stretching-based yoga, tai chi, or Pilates, in patients undergoing cancer treatment have mainly concentrated on their benefits in psychological aspects and treatment-related complications.^31–33^ In contrast, little is known as to whether mechanical stretching per se can elicit anti-tumor responses, despite emerging research on the counteracting effect of mechanical stretching on inflammation,^34^ which is a significant contributory factor to tumorigenesis.^35–37^ One of the main obstacles to understanding the basic mechanisms of stretching is the lack of animal *in vivo* models to mimic human self-stretching.^38^ An active stretching model established in several species (swine, rat and mouse) by our group has shown the anti-inflammatory effect of whole body stretching within subcutaneous tissues.^39–42^ More recently, our ongoing studies performed in a mouse orthotopic breast cancer model demonstrated that gentle active (tail lifting) and passive (under anesthesia) stretching inhibited breast tumor growth.^43, 44^ However, these stretching models may involve potential stresses caused by tail lifting and repeated anesthesia,^38, 45, 46^ emphasizing the necessity for a less stressful voluntary stretching model.

Therefore, the aim of the current study is to address this research need by developing and validating a new voluntary stretching mouse model to investigate the effects of stretching on mouse mammary tumor growth. Using food and water as motivators is a strategy employed by some types of animal exercise regimes.^47^ Based on this strategy, a previous study focusing on endurance exercise successfully established a voluntary stand-up rat model by progressively increasing the height of food in the cage.^48^ Accordingly, we hypothesized that voluntary stretching in mice could be achieved by using food and water as well as additional cage enrichment as motivators for stretching in their home cages.

## 2. Materials and methods

### 2.1 Animals

Six-week-old female FVB/NJ mice obtained from Jackson Laboratory were housed in the NIDCR Veterinary Resources Core under a 12-hour dark/light cycle. All animal procedures complied with the protocol approved by the Animal Care and Use Committee (ACUC). Animals were initially group-housed in regular home cages for a 1-week acclimation upon arrival, then pair-housed in behavior observation cages for 2 weeks. They had free access to food and water and were provided with paper pellet bedding (Teklad 7084, Envigo, Indianapolis, IN, US), which appeared dark in infrared light, throughout both the acclimation and behavior monitoring phases.

### 2.2 Experimental design

The experimental design of the current study includes the following steps (Fig. 1): 1) setting up hardware and software for behavior manipulation and tracking; 2) validating the effectiveness of the model in promoting voluntary stretching in long-term behavior tracking in a pilot study; 3) assessing anti-tumor effects of the new voluntary stretching model compared to voluntary running model; 4) using proteomics to explore the mechanisms underlying the effects of voluntary stretching relative to voluntary wheel running model.

**Fig. 1.**
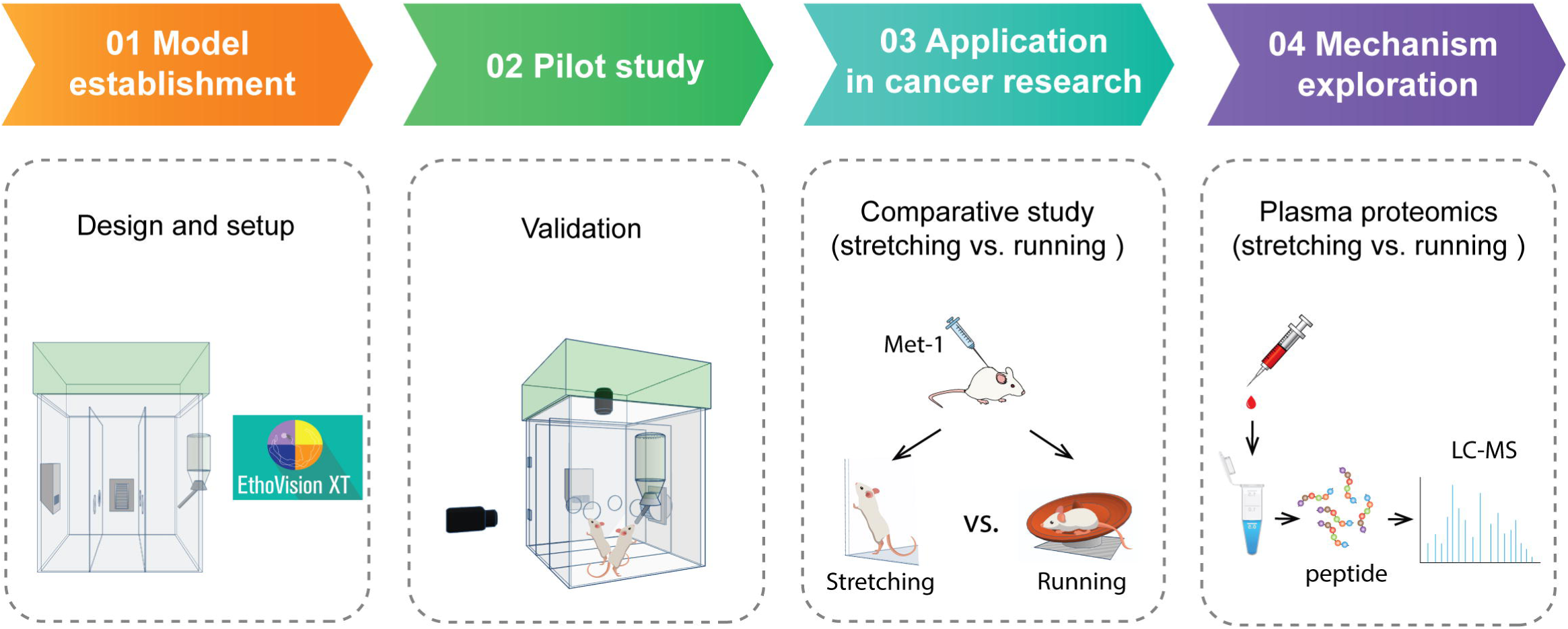
Schematic overview of the steps of experimental design of the current study. The study was undertaken in the workflow including behavior model establishment, testing the model in a pilot trial, application in a model of pathology, and exploration for relevant mechanisms.

### 2.3 Establishment of voluntary stretching model

#### 2.3.1 Behavior apparatus

We hypothesized that elevating the vertical distance to access water, food, and enrichment in animals’ home cages could increase their daily vertical activities. To test this, our design involved positioning the water bottle, food feeder and enrichment at an elevated location within the cage by using Noldus PhenoTyper 3000 customizable observation cages with its integrated video-based behavior recognition software, EthoVision XT 17 (Noldus, Wageningen, Netherlands) (Fig. 2A). To detect vertical stretching in the cage, a front-view infrared-sensitive camera (WAT-902H Supreme, Watec, Decatur, AL, US) was mounted at a height of 9 cm above the floor level, pointing towards an imaginary threshold line that indicates the event of a mouse’s vertical stretching. The preinstalled top-view infrared-sensitive camera is centrally positioned on the ceiling to capture horizontal activities (Fig. 2B). In the current study, the term “vertical stretching” denotes body elongation along the axis that is perpendicular to the floor, while “horizontal stretching” refers to elongation parallel to the floor.

**Fig. 2.**
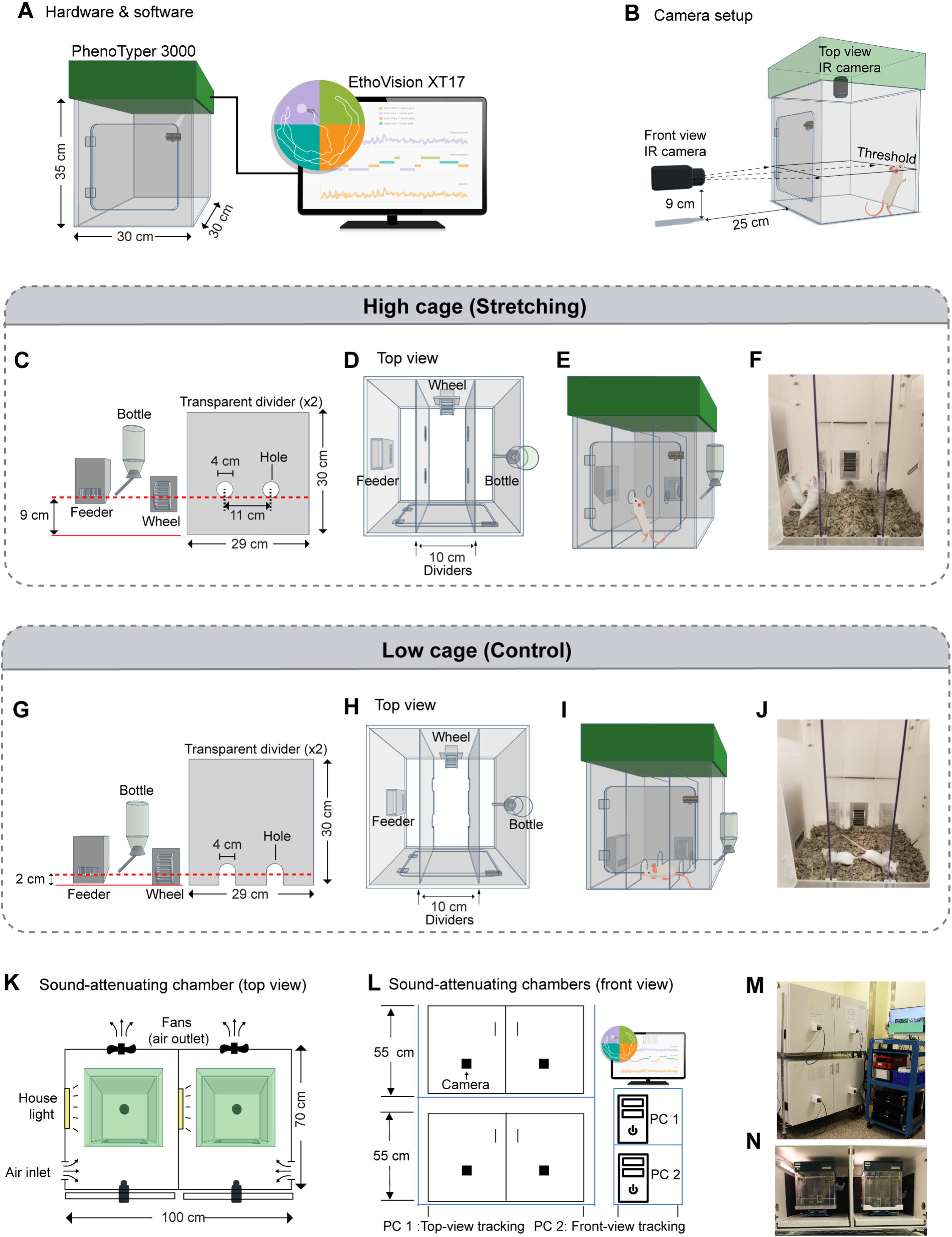
Setup for stretch-promoting and monitoring system. (**A**) Diagram showing the dimensions (30 cm W × 30 cm D × 35 cm H) of a Noldus PhenoTyper 3000 and Noldus behavior recording and analysis software, EthoVision XT17. (**B**) Schematic for the IR-sensitive camera setup for top-view and front-view video recording. The top-view camera is centrally positioned on the ceiling to capture the horizontal activities. The front-view camera, mounted at a height of 9 cm above the floor level, points towards an imaginary threshold line that indicates the event of a mouse’s vertical stretching. (**C-E**) Diagrams presenting the height and dimensions of cage accessories (C) in stretch-promoting high cage and the cage setup from top view (D) and front view (E). (**F**) A representative image showing two mice stretching in a high cage. (**G-I**) Cartoon showing the height and dimensions of cage accessories (G) in control low cage and the cage setup from top view (H) and front view (I). (**J**) An example picture showing two animals moving between cage compartments through dividers holes in a low cage. (**K**) Top view of the dimension and setup of a sound-attenuating chamber (100 cm W × 70 cm D × 55 cm H) that encloses two PhenoTypers. (**L**) Schematic presentation of front view of two sound-attenuating chambers stacked on a rack and the computer system for top-view tracking and front-view tracking. (**M, N**) Pictures showing the behavior system setup in the lab (M) and the interior look of a sound-attenuating chamber (N).

#### 2.3.2 Custom cage setup

Three custom cage features were designed to promote voluntary vertical stretching: 1) height-elevated food feeder and water bottle, ensuring animals extend their bodies vertically while accessing food and water; 2) height-elevated wall-mounted response wheel (ENV-113AS, Med Associate, Fairfax, VT, US) to enhance animal rearing posture; 3) two paralleled transparent cage dividers with height-elevated holes vertically placed between the feeder wall and bottle wall, inducing body elongation as mice shuttle between compartments to access food and water (Fig. 2C-E). Fig. 1F shows two FVB mice rearing up to reach an elevated feeder and a raised divider hole in the stretching-promoting cage (high cage). In contrast, the control cage (low cage) has these features at a lower position, allowing animals to access food and water without body elongation (Fig. 2G-I). Thus, the movements of low cage animals remained at the floor level (Fig. 2J). Both high and low cages were identical in size and shape. The terms “high cage” and “low cage” in this study only refer to the different locations of the feeder, bottle, and cage enrichment.

#### 2.3.3 Interior and exterior setup of sound-attenuating chambers

To minimize the interference of environmental factors on animal behaviors during 24/7 video tracking, two PhenoTyper home cages were enclosed in one customized sound-attenuating chamber (Maze Engineers, Skokie, IL, US) (Fig. 2 K). Each sound-attenuating chamber was equipped with two ventilation fans and LED lights controlled by a digital timer for the 12-hour light/12-hour dark cycle (light on at 7:00 AM and off at 7:00 PM). Top-view and front-view cameras connected to two separate workstations allowed for independent recording of the two perspectives (Fig. 2L). Fig. 2M and 1N show the exterior and interior setup of the behavior monitoring system.

#### 2.3.4 Behavior analysis

Vertical and horizontal stretching were automatically analyzed by the software Noldus EthoVision XT 17. For a full description of how to quantify vertical stretching and horizontal movement in EthoVision, see Supplementary methods, “Vertical stretching quantification and verification” and “Horizontal movement analysis”. Briefly, vertical stretching was tracked through a strategy of detecting the event of the body ends crossing the designated threshold lines in both high cage and low cages. The frequency and time spent on the events were recorded as the vertical stretching frequency and duration. As for horizontal movement, we tracked horizontal stretching and horizontal locomotion, respectively using the body-shape-based “Body elongation” and body-point-based “Distance moved” features in the software. Note that during long-term video tracking, identity swaps occurred frequently between the pair-house animals with identical appearance. However, EthoVision has technical limitations in automatically correcting these swaps. Therefore, for the automated behavior analysis in this study, we used cage-based average stretching frequency/duration to reflect the behavior performance of the pair-housed animals in each cage.

### 2.4 Grip strength test

To access the effects of stretching on the overall muscle strength in this voluntary stretching model, grip strength test was performed at the end of the 2-week monitoring in the pilot study. Details of the procedure is described in the Supplementary methods, “Grip strength test”.

### 2.5 Orthotopic mammary tumor model

A well-established mammary cancer mouse model in our lab was utilized to evaluate the efficacy of the voluntary stretching model.^44^ Briefly, a mouse mammary tumor cell line, MMTV-PyMT (Met-1), was orthotopically injected, at the amount of 2×10^4^ tumor cells in 100□µl mix prepared: 60% DMEM and 40% Geltrex (Thermo Fisher Scientific, Waltham, MA, US), into FVB mice bilateral third mammary fat pads. The day after injection, animals were transferred from regular home cages to PhenoTypers to undergo 2-week behavior monitoring. The growth of the tumor was measured weekly by caliper and ultrasonography while the animals were under anesthesia, as previously described.^43, 44^

### 2.6 Voluntary wheel running

To compare the effectiveness of voluntary stretching with another exercise model for suppressing mammary tumors, we conducted a comparative study using voluntary wheel running, a common preclinical aerobic exercise model in cancer research. To ensure environmental conditions were similar for both exercises, running wheels were placed in PhenoTyper low cages, with feeders and bottles set at the floor level to minimize the animals’ vertical stretching. The day after receiving tumor injection, mice were pair-housed in PhenoTyper cages with two wireless mouse running wheels (ENV-047, Med Associate, Fairfax, VT, US) in each cage. The running group had 24/7 access to wheels for two weeks, while the wheels for the sedentary group remained locked throughout the experiment. The wheel activity was recorded by the Wheel Manager Software (Med Associate, Fairfax, VT, US). Similar to the voluntary stretching model, running performance was presented as cage-based average revolution and distance, given that the pair-housed mice were allowed to access both running wheels in the same cage.

### 2.7 Body composition by dual energy X-ray absorption (DEXA) scan

The impact of stretching on body composition was assessed via DEXA scanning. Detailed procedure is described in the Supplementary methods, “DEXA scan”.

### 2.8 Plasma proteomics and bioinformatic analysis

Plasma proteomics analysis was conducted on the blood samples collected upon the endpoint of the voluntary stretching and voluntary running experiments. A detailed description of sample preparation and liquid chromatography-mass spectrometry (LC-MS) is provided in the Supplementary methods, “Sample preparation and LC-MS Assay”. The proteomics data was visualized using SRplot,^49^ an online tool available at https://www.bioinformatics.com.cn/en. Gene Ontology (GO) enrichment analysis was performed by using the Database for Annotation, Visualization and Integrated Discovery (DAVID 2021).^50^

### 2.9 Statistical analysis

Two sample t-tests were used to compare cage-based outcomes that were measured at a single timepoint. When applicable, the Satterthwaite approximation was used to estimate the degrees of freedom when there was evidence that group variances were not homogeneous. Mixed model repeated measured analyses were used to compare groups on cage-based outcomes measured at multiple timepoints (i.e. weeks). Group comparisons on animal-based outcomes were performed based on a nested mixed model (i.e. animal nested within cage). Nested mixed model repeated measures analyses were used to compare groups on animal-based outcomes measured at multiple time points. Simple effects (i.e. group comparisons within timepoint) were based on partial *F*-tests. Tumor volume was log transformed prior to analysis to account for heterogenic variances across time points. Geometric means and associated standard errors are presented when data was log-transformed. Statistical analyses were performed using SAS statistical software (SAS Institute, Cary, NC, US).

## 3. Results

### 3.1 Behavior performance and physiological response in long-term observation

Vertical stretching was measured using a software-based automated strategy to detect the event of the mouse body end (nose or tail) entering the designated threshold zone, as depicted in Supplementary methods and Fig. S1A-L. In a pilot study where we tested the algorithm, we compared manual and automated scoring of vertical stretching in a 15-minute video segment to verify the accuracy of software-based automated analysis (Fig. S1M-O). Then, we extended the length of observation to two weeks for continuous 24/7 behavior monitoring. During the 2-week pilot testing period, high cage mice were observed to vertically cross the threshold when accessing food, water, response wheel, and divider holes positioned at an elevated level (Fig. 3A-C and Supplementary Movie 1), while low cage mice predominantly engaged in these activities at the floor level (Fig. 3D-F). Automated analysis revealed that high cage mice showed at least 2-fold increase in the frequency and duration of vertical stretching compared to low cage mice (Fig. 3G). In addition, the heat-map plots in Fig. 3H visualized the hotspots that animals frequently visited in the PhenoTypers, showing that high cage mice increased their vertical movement and spent more time at the locations of elevated feeders and dividers with high holes, as opposed to their low cage counterparts. Moreover, a representative time-event plot in Fig. 3I highlighted the frequency/duration difference of the entry of animals’ body ends in the “threshold zone” between high cage and low cage animals during active hours.

**Fig. 3.**
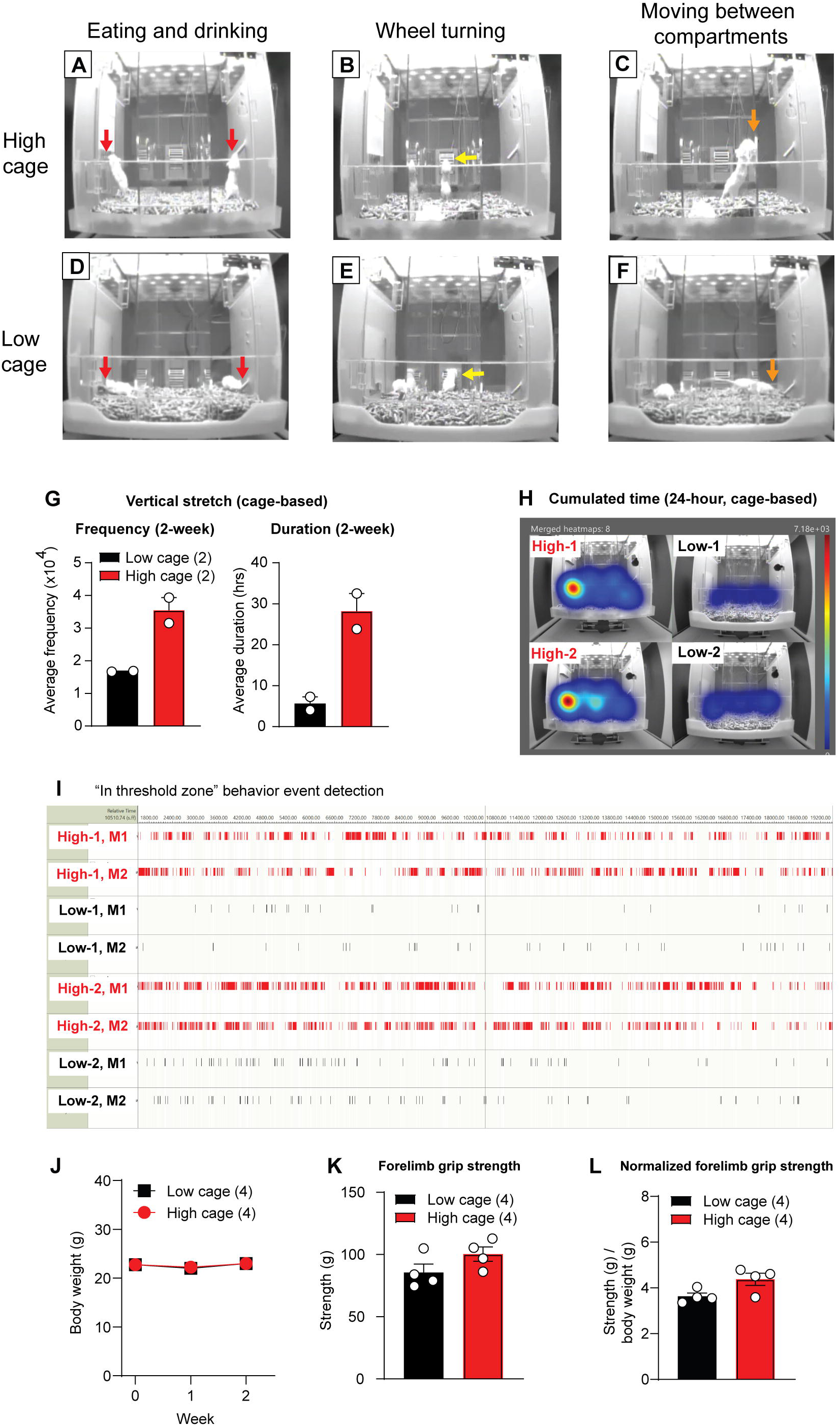
High cage setting enhances vertical stretching in mice in pilot study. (**A-C**) Front-view camera images showing the representative above-threshold vertical stretching which were exhibited during eating and drinking (A), wheel turning (B) and crossing holes (C) in the high cage animals. The typical behaviors were denoted by the arrows. The threshold of vertical stretch was visualized by a white string attached the front wall. **(D-F)** Images showing that, in the low cage, eating, drinking (D), wheel turning (E), and hole crossing (F) behaviors predominantly occurred below the threshold. The arrows denote the animals displaying these behaviors. (**G**) High cage animals versus low cage animals displayed an increase in frequency and duration in a two-week behavior tracking period (cage-based, n=2 per group, each dot denotes one cage). Cage-based strategy was used for presenting all automated analysis results (see Material and Methods). (**H**) A heatmap visualization demonstrating the hotspot visited by high cage and low cage mice based on 24-hour video footage. (**I**) A representative time-event plot illustrating the frequency and duration of the vertical stretching in high cages and low cages during active hours. (**J**) No difference in body weight was observed between high-cage and low-cage animals within the 2-week behavior tracking (animal-based, n=4 per group, n represents the animal number in each group, *p=*0.97). (**K, L**) High-cage animals versus low-cage animals exhibited a slight, but not significant, increase in forelimb grip strength (K, n=4 per group, each dots represents an animal, *p=*0.36) and forelimb grip strength normalized to body weight (L, n=4 per group, each dots represents an animal, *p*=0.051) at the end of the two-week behavior monitoring. All values are represented as mean ± SEM. Statistics: (J): *F*-test from nested repeated measures mixed model; (K) and (L): *F*-test from nested mixed model.

Interestingly, in the extended observation phase (> 3 weeks) of the pilot study, horizontal behavior video tracking (Fig. S2A-R) also revealed that some older animals (>12-week-old) with larger body sizes stretched horizontally while not in contact with the floor. This behavior was promoted by the elevated enrichment in the high cage, especially when high cage animals used elevated divider holes to travel between different compartments, as larger animals were able to stretch between the divider holes, the distance of which was close to their body length. (Fig. S2I-P and Supplementary Movie 2). Taken together, these data suggest that using water, food, and cage enrichments as reinforcers can effectively facilitate animal self-stretching behavior over the long term.

To track whether vertical stretching would impact animals’ energy expenditure, we monitored their body weight each week during the 2-week behavior testing. No significant difference was found in body weight between the high cage and low cage animals (Fig. 3J). However, we observed a minor, not statistically significant trend in increased grip strength (Fig. 3K) and normalized grip strength (Fig. 3L) in the high cage animals at the end of the 2-week experiment, suggesting that the voluntary stretching may impact overall muscle strength.

### 3.2 Application of the new voluntary stretching model in breast cancer research

We further tested the potential of the new voluntary stretching model as an exercise intervention to reduce the growth of mammary tumors, a pathological condition in which active and passive stretching have shown positive outcomes in our previous studies.^43, 44^ We sought to determine whether the voluntary stretching model is similarly effective as the voluntary running model, one of the most commonly used aerobic exercise models to investigate the roles of physical activity on mammary cancer. To achieve this, we performed a comparative study including two types of physical activities (voluntary stretching and voluntary running) relative to their respective experimental controls (low cage animals for voluntary stretching and animals provided with locked running wheels—referred to as sedentary control—for voluntary running).

As shown in the experimental timelines (Fig. 4A and Fig. 4I), both methods involved orthotopic injection of the Met-1 tumor cells into the dorsal fat pads of FBV mice. We then separately examined the anti-tumor effects of voluntary stretching and wheel running over a 2-week observation period. In the voluntary stretching model, high cage mice had a greater frequency and duration of vertical stretching compared to low cage mice (Fig. 4D) in each single week of observation. This resulted in a significant reduction in the tumor growth rate, as measured by 3D-ultrasound and caliper (Fig. 4B and C). In the voluntary wheel running model, running mice had an increased wheel revolution and distance over time, compared with sedentary controls (Fig. 4L), as well as a significant reduction in tumor growth measured by 3D-ultrasound and caliper (Fig. 4J and K). The tumor-suppressing effect observed in the high cage animals was mainly attributed to vertical stretching, as there was no difference in the horizontal stretching between the high and low cage mice within the 2-week observation (Fig. S2S-U). In addition, although there was a slightly greater total distance traveled in the high vs. low cage animals (Fig. S2V), this difference was much smaller compared to the difference in running distance between voluntary wheel running and sedentary control (Fig. 4L). Thus, it is unlikely that the slightly increased horizontal locomotion in high cage animals was responsible for the tumor inhibition observed in the voluntary stretching model.

**Fig. 4.**
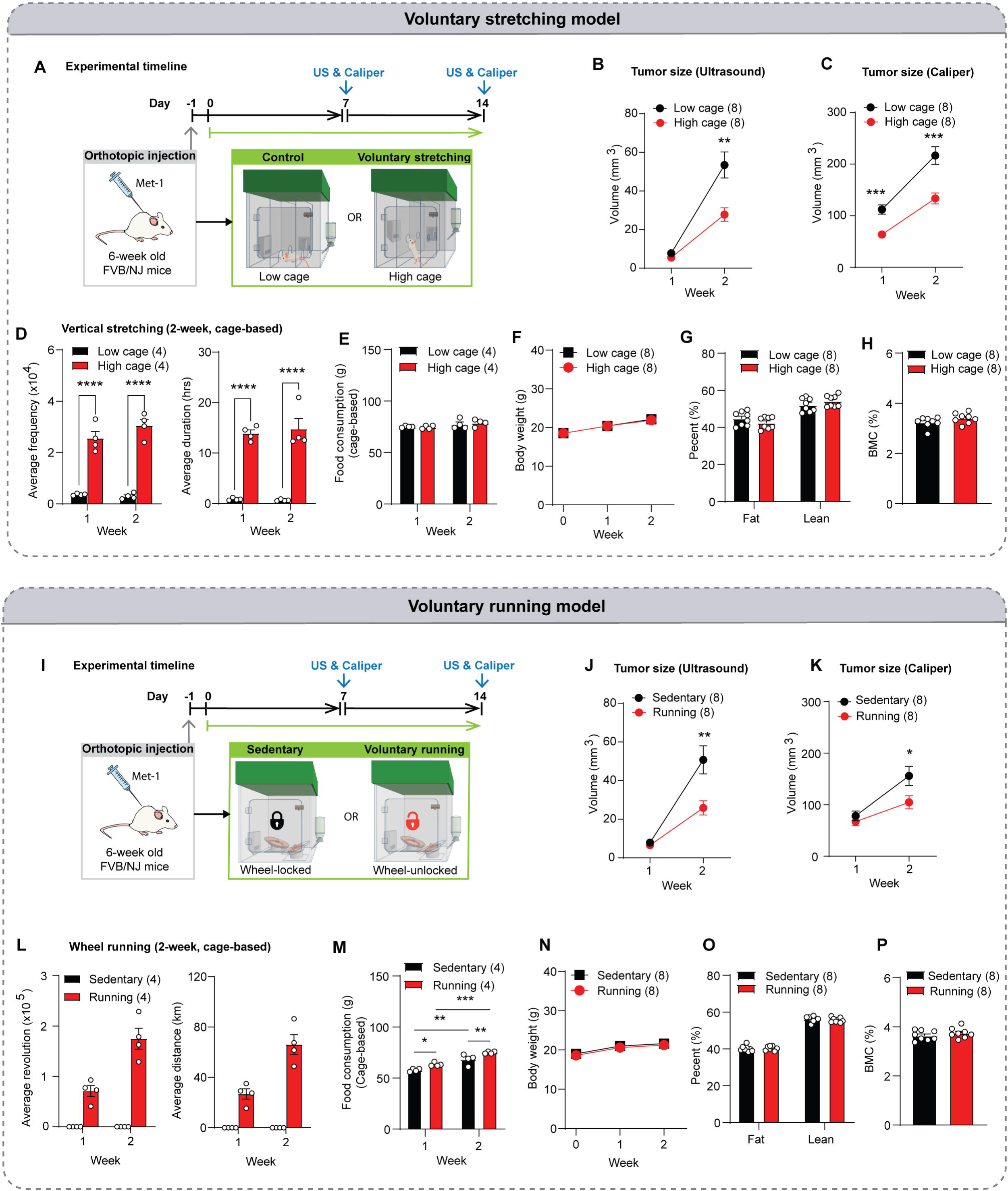
Effects of voluntary stretching and voluntary running on breast tumor growth. **TOP PANEL: Voluntary stretching model.** (**A**) Experimental timeline for evaluating the effects of voluntary stretching (high cage vs. low cage) on orthotopic mammary tumor growth. (**B, C**) Vertical stretching significantly inhibited tumor growth (animal-based, n=8/group), measured by ultrasound (B, week1: *p*=0.081; week 2: *p=*0.005) and caliper (C, week 1 and 2: *p*<0.001). (**D**) High cage animals had a significant increase in average stretching frequency and duration (cage-based, n=4/group; stretching frequency/duration at week 1: *p*<0.0001; stretching frequency/duration at week 2: *p*<0.0001). Cage-based strategy was used for presenting all automated analysis results. (**E**) Vertical stretching did not significantly change weekly food intake (cage-based, n=4/group) in high cage versus low cage animals (week 1: *p=*0.85; week 2: *p=*0.56). Cage-based strategy was used to show the average weekly food consumption per animal due to the pair-housed setup. Average food intake= (Initial food weight-Remaining food weight)/2. (**F**) Vertical stretching did not alter animal body weight (animal-based, n=8/group, *p=*0.94). (**G, H**) Body composition analysis (animal-based, n=8/group) showing that 2-week vertical stretching did not alter fat percent, lean percent (G, fat percent: *p=*0.39; lean percent: *p=*0.41), or bone mineral content (BMC) (H, *p=*0.18l). **BOTTOM PANEL: Voluntary running model.** (**I**) Experimental timeline for assessing the anti-tumor effect of voluntary running. (**J, K**) Tumor growth (animal-based, n=8/group) was significantly suppressed by voluntary running (compared with sedentary) as measured by caliper (J, week 1: *p*=0.38; week 2: *p*=0.003) and to a lesser extent by ultrasound (K, week 1: *p*=0.37; week 2: *p*=0.028). (**L**) Running animals showed a progressive increase in weekly running revolution and distance over time. Cage-based strategy was used for presenting average running performance. (**M**) Running and sedentary animals displayed significantly increased weekly food intake (cage-based, n=4/group), with both showing gradually increased food consumption over the weeks (week 1: *p=*0.036; week 2: *p=*0.015; sedentary: week 1 vs. week 2, *p=*0.001; running: week 1 vs. week 2, *p*<0.001). (**N**) 2-week voluntary running did not alter body weight (animal-based, n=8/group) in running versus sedentary animals (*p=*0.57). (**O, P**) 2-week voluntary running did not change body composition (animal-based, n=8/group), including fat and lean percent (O, fat percent: *p=*0.39; lean percent: *p=*0.42; P, BMC: *p=*0.35). The tumor volume data is presented as the geometric mean ± SEM, while all other data is presented as the arithmetic mean ± SEM. Statistics: (B) to (E), (J), (K), and (M): partial *F*-test from nested repeated measures mixed model; (F) and (N): *F*-test from nested repeated measures mixed model; (G), (H), (O), and (P): *F*-test from nested mixed model. [\**p* < 0.05, \*\**p* < 0.01, \*\*\**p* < 0.001, \*\*\*\**p* < 0.0001].

Given that energy balance is linked to cancer development and progression,^51–53^ we also monitored whether animals’ cumulative food intake, body weight, and composition were altered by exercise-induced energy expenditure. In the voluntary stretching model, we found that the animals’ cumulative food intake remained unchanged regardless of their activity levels in high or low cage environments (Fig. 4E). In the voluntary wheel running model, food intake was not only increased in the running group compared to the sedentary control but also over time (Fig. 4M). For body weight, all groups displayed gradual gain over time. Animals with higher physical activities in both models did not show a significant change in body weight compared to their less active controls during the 2-week observation period (Fig. 4F and N). Considering that obesity is associated with poor outcomes in early breast cancer^54^ and exercise promotes positive changes in body composition, we further evaluated the post-mortem body composition at the end points of the experiments. High cage animals showed only a slight, but not significant reduction in fat percent and a minor increase in lean percent as well as bone mineral content (BMC) compared with the low cage animals (Fig. 4G and H, low cage vs. high cage: fat percent: 44.7% vs. 42.4%; lean percent: 52.0% vs. 54.2%; BMC: 3.2% vs. 3.4%). In contrast, in the voluntary running model, the body composition of running animals was not significantly different from that of sedentary animals (Fig. 4O and P, sedentary vs. running: fat percent: 40.2% vs. 40.5%; lean percent: 56.2% vs. 55.7%; BMC: sedentary vs. running, 3.6% vs. 3.8%). However, our observation was not extended to assess the longer-term effects of voluntary stretching and voluntary wheel running, which may impact energy balance and body metabolism differently.

### 3.3 Voluntary stretching versus voluntary running differently impacts plasma proteomics profiling

To compare how voluntary stretching and voluntary wheel running influence biological processes and the mechanisms behind their anti-tumor effects, we employed LC-MS-based quantitative proteomic approach to systemically explore the plasma proteome changes after exercise. Our findings revealed notable disparities in protein expression profiles between the two voluntary exercise modalities. Specifically, we identified a total of 1394 and 1082 proteins, respectively, in voluntary stretching and voluntary wheel running models, with 864 proteins common in both models (Fig. 5A). Upon setting the significance threshold to *p*-value<0.05, |Fold change|>1.5, we found that the high cage group in voluntary stretching model exhibited an upregulation of 273 plasma proteins and a downregulation of 36 plasma proteins compared to the low cage group (Fig. 5B). In the voluntary wheel running model, 47 upregulated proteins and 82 downregulated proteins were observed when comparing the running group to the sedentary group (Fig. 5C). An identified protein, A0A3B01TG8 (Adiponectin), showed a 12.4-fold increase (log2 fold change=3.63) in voluntary stretching animals and only a 1.3 -fold increase (log2 fold change=0.38) in voluntary wheel running group. It was marked in both volcano plots as an example of a protein that may be differentially regulated in the two different models (see Discussion). Venn diagrams in Fig. 5D and E further illustrated that only 6 proteins were commonly upregulated in both the voluntary stretching and voluntary wheel running (Fig. 5D), and 4 proteins were downregulated in both (Fig. 5E). To better understand the functional implications of these differentially expressed proteins, we performed Gene Ontology (GO) enrichment analysis on the identified protein sets. A side-by-side comparison of the top five GO terms revealed that the proteins altered by voluntary stretching and voluntary wheel running were associated with distinct functions across three GO categories: biological process (Fig. 5F and I), cellular components (Fig. 5G and J), and molecular function (Fig. 5H and K). These data offer a comprehensive perspective on the prominent difference in cellular and molecular mechanisms between voluntary stretching and voluntary wheel running exercise models.

**Fig. 5.**
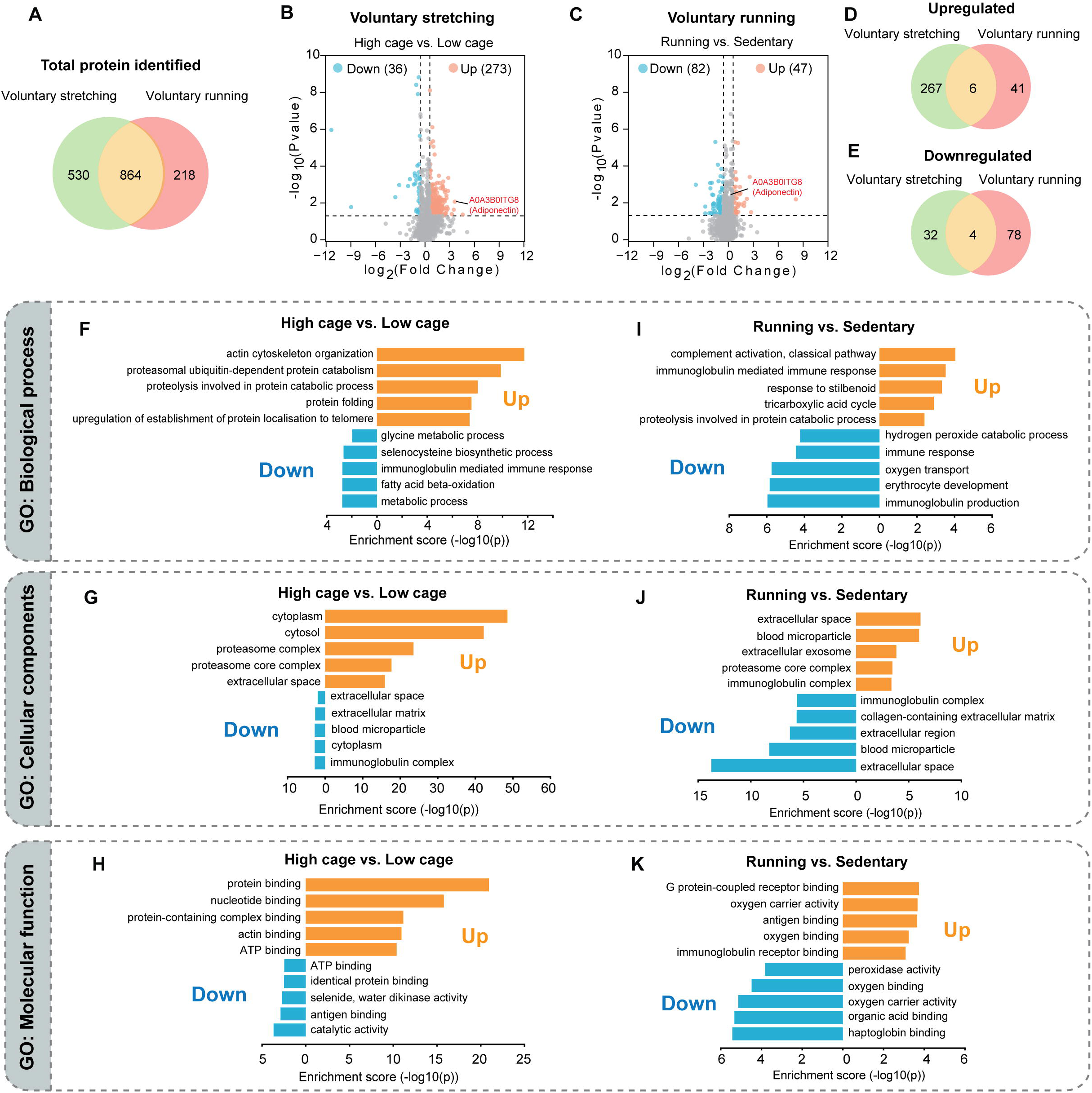
Voluntary stretching vs. voluntary running impacts differently on plasma protein profiles and the pathways involved. (**A**) A Venn diagram showing the overlap between the total number of proteins identified in the voluntary stretching model and the voluntary wheel running model. (**B, C**) Volcano plots displaying the distribution of the all proteins identified from the voluntary stretching model (B) and the voluntary wheel running models (C) with log_2_ fold change values plotted against their associated negative log_10_ p-values. Blue dots and orange dots respectively represent individual proteins with significant increase and reduction (*p*<0.05, absolute fold change |FC|>1.5). An identified protein, A0A3B0ITG8 (Adiponectin), was labeled in both volcano plots as an example of expression comparison. (**D, E**) Venn diagrams showing overlaps of significantly upregulated (D) and downregulated (E) proteins between the voluntary stretching model and the voluntary wheel running model (*p*<0.05, absolute fold change |FC|>1.5). (**F-K**) Gene ontology (GO) enrichment analysis of voluntary stretching (F-H) and voluntary wheel running (I-K) models showing top 5 (ranked by *p*-value) GO terms that upregulated (orange) and downregulated (blue) proteins are over-represented in three categories (biological process, cellular components, molecular function). All groups contained plasma collected from eight animals (n=8), except for the high cage group (n=7), in which the plasma volume in one sample was insufficient to run the assay.

## 4. Discussion

### 4.1 The first voluntary stretching exercise model with minimal stressors

Our study developed the first mouse voluntary stretching model, which was effective in promoting mouse vertical stretching over a two-week observation period. The results from this voluntary stretching model are consistent with our previous major findings on the impact of human-handling active stretching on mammary tumor growth.^43, 44^

Animal behavior models are essential tools in biomedical research as they facilitate the investigation of the intricate interplay between behavior and physiological mechanisms. Aerobic, resistance, and flexibility exercises are three major physical exercises recommended by physical activity guidelines.^55^ Compared to the modeling of human aerobic and resistance exercises,^47, 56, 57^ the study on a full-body stretching-based flexibility exercise animal model still lags behind.^38^ Existing animal models of stretching mainly focus on utilizing stretch devices to induce localized mechanical tension in the muscles of anesthetized animals to investigate muscle hypertrophy^58, 59^ and injuries.^38^ Few animal studies simulate the whole-body myofascial stretching that is usually performed in human yoga practice.^38^ Our previous tail-lifting active stretching model has the advantage of facilitating full-body extension, but it is somewhat similar to the procedures used in the tail suspension test^60^ and the hindlimb suspension model,^61^ which raised the concerns about the potential mental and physical stress.^62, 63^ Minimizing the potential stress responses in animals is a fundamental consideration when designing an animal exercise protocol.^47^ The new voluntary stretching model addressed this goal by eliciting animal rearing, an innate behavior displayed in rodents when they engage in exploring the environment. Similar to the prevalent voluntary wheel running model, which was driven by intrinsic exercise reward, voluntary stretching was motivated by food and water rewards and environmental enrichment, reducing the adverse health impact and confounding outcomes produced by forced exercise.^64^

### 4.2 Voluntary stretching as a promising low-intensity exercise modality for cancer research

Besides being a less stressful stretching model, the voluntary stretching model could serve as a lower-intensity exercise alternative to voluntary running, its vigorous-intensity aerobic counterpart. In the experiment comparing the tumor-suppressing effects of the two voluntary exercise models, voluntary stretching, despite being a more moderate form of exercise, showed comparable results to voluntary running in reducing mammary tumor growth in the same breast cancer model.

Regarding exercise intensity, clinical trials have shown a trend in introducing high-intensity interval training (HIIT), a workout that combines vigorous aerobic and strength training, to breast cancer survivors.^65, 66^ However, challenges such as the intensity of HIIT exercises, requirement for professional supervision, and access to exercise facilities have affected patient adherence to the program, resulting in the intervention being primarily lab-based rather than home-based.^65, 67^ Moreover, limited information and investigation are available regarding whether high-intensity exercise can slow cancer progression and reduce metastatic dissemination in breast cancer patients. In the preclinical context, while many have explored the positive effects of aerobic exercise on mammary tumor-bearing animals, there are still controversies and questions surrounding these findings. Forced swimming and forced treadmill running in aerobic exercise model have been reported to accelerate mammary tumor growth and increase metastatic spread in lungs.^16–18, 20, 68, 69^ Even the least stressful aerobic exercise model, voluntary wheel running, has raised concern over its risk of increasing mammary cancer lung metastasis in a 4T1 orthotopic mammary cancer BALB/C mouse model,^19^ which is a less-investigated aspect of the exercise-cancer hypothesis. Conversely, another group demonstrated that wheel running did not affect pulmonary metastasis in the same type of mammary cancer model. However, the result may be misinterpreted because the mean value of lung metastasis in the sedentary group was overestimated due to a few outliers with higher values. In fact, the median of lung metastases in the running group was much higher than that in the sedentary group.^24^ Our current study did not compare the lung metastasis between voluntary stretching and voluntary running models due to the lack of visible metastatic lung nodules at the experiment endpoint in week two. However, one of our unpublished preliminary studies, which lasted three weeks, indicated that active stretching did not increase visible metastatic tumor nodules on the lung surface compared to the control group without stretching, as examined by the Indian ink inflation staining method.^44^ These findings highlight the importance of careful assessment of the potential for different exercise models to promote metastasis in our future studies.

Additionally, it is noteworthy that low-intensity voluntary stretching can also effectively activate certain protective pathways involved in cancer prognosis despite its low-intensity feature. For example, one of the overrepresented identified proteins in the voluntary stretching group, A0A3B0ITG8 (Adiponectin) (labeled in Fig. 5B), is a hormone produced and secreted by adipose tissue. It has been increasingly studied for its role in breast cancer, with evidence suggesting it may have protective effects against tumor development and progression, through its ability to regulate fatty acid metabolic remodeling and reduce inflammation.^70, 71^ Low serum adiponectin levels have been reported to be associated with obesity and aggressiveness of postmenopausal breast tumor, suggesting that adiponectin levels could be used as a biomarker to predict breast cancer survival.^72, 73^ This may support our findings in proteomics analysis where we observed a significant increase in adiponectin levels in the voluntary stretching group compared to only a marginal increase in the voluntary running group, suggesting that stretching may specifically activate adiponectin to play a role in its anti-tumor effects (Fig. 5B and C). The bioinformatic analysis of proteomics data in our study not only provides a comprehensive view of the proteome changes induced by stretching but also delineates the directions of future mechanistic investigations into its anti-cancer effect.

Together, given that the intensity of physical activity is an important aspect to evaluate before prescribing exercise interventions to cancer patients, the low-intensity and anti-tumor features observed in the voluntary stretching model suggest the possibility of developing a home-based, low-risk preventive intervention for breast cancer survivors. The mechanistic insights gained from our voluntary stretch model may pave the way for its clinical application and provide valuable preclinical evidence to guide clinical decision-making.

### 4.3 More potential applications of voluntary stretching model

Stretching is a key component of yoga, a popular mind-body exercise known for its multifaceted benefits across various body systems, including reducing chronic musculoskeletal pain, preventing cardiovascular diseases, alleviating mental stress, enhancing the immune system and mitigating inflammation.^74–77^ However, it remains largely unexplored regarding the interface and connection between the mechanical signaling in the general connective tissue (e.g. muscle, bone and fascia) and physiological responses in other body systems. Additionally, while the voluntary wheel running model is used extensively to study the health benefits of exercise in various research fields, its benefits may be limited to mechanisms associated with vigorous aerobic exercise. The proteomics analysis of the plasma proteins of voluntary stretching and voluntary wheel running models in our current study also shows that the mechanisms behind their physiological responses are model-specific. To date, no prior research has addressed how yoga benefits multiple body systems using an animal stretching behavior model, indicating this novel voluntary stretching model has the potential to be a research tool to open new avenues for mechanistic studies on the health benefits of stretching-based exercises across various research fields and fill existing knowledge gaps in these areas.

### 4.4 Limitations

Although the present study offers a novel method to lay the groundwork for future research in voluntary stretching, several limitations need to be noted. Firstly, unlike the previous active and passive stretch models, the parameters/dose of voluntary stretching (intensity, frequency, timing, duration) are uncontrollable in the current system, similar to the limitation of voluntary wheel running versus forced treadmill running. This limitation may restrict the applicability of the voluntary stretching model to studies requiring controllable levels of exercise. Secondly, since animals automatically conduct voluntary stretching, the vertical stretching recorded in the system only represents the general whole-body stretch rather than reflects specific posture-related stretches in human yoga. Alternatively, the passive stretch model could be an option if the stretch direction relative to the body axis is the research focus. Thirdly, one issue that remains unaddressed is the identification of individual animal stretch behavior during multi-subject tracking. In our study, two mice were housed in each observation cage for social enrichment and animal welfare. The video tracking software EthoVision encountered an issue of subject identity swaps when two markerless animals with identical appearance touched each other during long-term video tracking. In fact, maintaining identity remains one of the most challenging aspects of tracking multiple freely moving animals with identical appearances over long-term continuous recording.^78, 79^ Consequently, we used the cage rather than the individual animal as the experimental unit for behavior quantification. The mean value from the two pair-housed animals was used to compare behavior performance between the experimental and control cages. Therefore, our study is unable to study the correlation of the tumor size to the behavior of each individual animal. One potential solution involves using machine learning to integrate imaging processing from the camera with a radio-frequency identification (RFID) signal could be a promising solution to correct the identity swaps.^80^

## 5. Conclusion

This study presents a new voluntary stretching model using food, water, and enrichment in the home cage as motivators for stretching. Using this model, we demonstrate that voluntary stretching inhibits cancer growth in the MET-1 orthotopic mammary tumor FVB mouse model. Although still in its nascent stage, this new model will potentially advance the mechanistic study of the health benefits of stretching in cancer research and a wide range of biomedical research areas.

## Acknowledgments

We thank NIDCR Veterinary Resources Core for their assistance in setting up the behavior tracking system and animal caring and Dr. Zheng-Xiong Xi from NIDA for help in mouse wheel running behavior tracking.

## Author contributions

YH participated in the design of the study, data collection, data analysis and writing; LB participated in the design of the study, data collection, data analysis and writing; YW contributed to data collection, data analysis and writing; GJB contributed to data analysis and writing. HML participated to in the design of the study, supervision, funding acquisition, reviewing and editing. All authors have read and approved the final version of the manuscript, and agree with the order of presentation of the authors.

## Abbreviation

BMC: Bone mineral content
DAVID: Database for Annotation, Visualization and Integrated Discovery
DEXA: Dual energy X-ray absorption
GO: Gene ontology
LC-MS: Liquid chromatography–mass spectrometry
MMTV-PyMT: Mouse mammary tumor virus-polyoma middle tumor-antigen
NIDCR: National Institute of Dental and Craniofacial Research

## Supplementary materials

## Supplementary methods

### Vertical stretching quantification and verification

Vertical stretching was tracked by the front-view camera and quantified through a strategy of detecting the event of the nose or tail crossing the designated threshold lines. To ensure the horizontal field of view (HFOV) of the front-view camera was overlaid with a threshold plane, two white strings were affixed at the height of the threshold (9 cm above the PhenoTyper floor) on both front and rear walls. Precise alignment was achieved by adjusting the height using a lab jack placed under the PhenoTyper to ensure the exact overlay of the front and rear threshold lines in the front view.

We first performed a pilot study with a smaller group size and shorter observation time to establish the behavior analysis algorithm and verification. The algorithm for analyzing vertical stretching is generally based on the “in zone” analysis profile in EthoVision XT 17, which counts the frequency the animal visited the defined zone and the total time spent in the zone. For the low cage, we defined a zone above the threshold line as a “threshold zone” in the front-view arena (Fig. S1A and B) and used “nose-tail base detection” feature in EthoVision XT 17 to track the “threshold zone” entries of animals’ body points. The vertical movement of animals in the low cage is primarily upward stretching against cage walls (Fig. S1C). Despite the body/tail point swapping during tracking (e.g. nose-point detected as tail-based and vice versa), the accuracy of results was not affected since the determination of the vertical elongation state only depended on the entry of either of the body ends, regardless of which end of the body was involved. Fig. S1D illustrated the common algorithm for calculating the upward vertical stretching in both cages: if either the nose point or tail point entered the zone, signifying the condition “in threshold zone” =true, the frequency of the event and duration of the state were recorded. For example, once the nose point of an animal enters the “threshold zone”, the software identifies the event, records the frequency as one, and the time of the presence of the body end in the zone, reflecting the upward vertical stretching frequency and duration. Conversely, the event was not counted in the condition that the body points were outside the threshold zone.

An additional step was included in the strategy of quantifying vertical stretching in the high cage animals, as compared to their low cage counterparts, animals in the high cage exhibited additional downward stretching when they moved between three compartments through high divider holes (Fig. S1G). The algorithm in Fig. S1D does not count the subsequent downward stretching after animal enters the “threshold zone”, as the “in zone” feature only counts once when either of the body ends enters the zone, without counting the event of the body exiting the zone. As shown in Fig. S1E-G, the hole crossing behavior consisted of three phases: moving towards the hole (E, upward stretching), nose and tail being within the hole zone (F, no stretch), and moving away from the hole (G, downward stretching). Therefore, besides the “threshold zone” (Fig. S1H and I), an additional “high hole zone” was introduced in the arena to specifically track the downward stretching during hole crossing (Fig. S1J). Fig. S1K shows the flowchart to quantify the downward stretching around the “high hone zone”. When the condition “Nose point AND tail base in the hole zone” is met, it predicts the subsequent downward stretching. The frequency of downward stretching matches the frequency of “nose point AND tail based in hole zone”. Thus, the total vertical stretching frequency in the high cage is the sum of frequency of “Nose point OR tail base” in threshold zone and frequency of “Nose point AND tail base” in hole zone (Fig. S1L). The duration of vertical stretching in the high cage is calculated by subtracting “in hole zone duration” from “in threshold zone duration”, as both the nose point and tail base being in the hole zone reflects a state that the animal’s posture transiently returned to “no stretch” (Fig. S1F).

To validate the software-based automated detection of vertical stretching, manual scoring of vertical stretching was also performed on the same video clip by the experimenter using the manual scoring feature in EthoVision XT 17. A 15-minute video footage was chosen, during which both high and low cages were recorded simultaneously, and the animals in a high cage exhibited active vertical stretching. Each of the pair-housed animals was assigned a specific key on the computer keyboard. The key was pressed each time when the observer judged that the animal was initiating stretching and completing the behavior. The length between the start and stop time points was recorded as duration. Compared to manual scoring, identity swaps in the pair-housed animals are inevitable in the automated analysis due to the current technical limitation of Noldus EthoVision XT. Thus, we used a cage instead of an individual animal as an experimental unit for the automated behavior analysis in the current study. The cage-based mean value of stretching frequency and duration from the two pair-housed animals was used to compare the behavioral performance between high cage and low cage.

### Horizontal movement analysis

Horizontal stretching was recorded by the top-view camera and detected by the user defined threshold based on the elongation percentage of the animal’s body. Horizontal stretching posture was specified as ≥ 90% of the maximum body elongation length. The animal’s total horizontal distance moved was monitored and measured by tracking the distance traveled by the center point of an animal. Due to the challenge of maintaining identity recognition in multi-subject tracking in behavior monitoring software, the cage-based strategy was also used to calculate the average horizontal stretching frequency/duration and locomotion for the two animals in the same cage.

### Grip strength test

An animal was allowed to grasp a metal T-bar mounted on a grip strength test meter (GT3, Bioseb, Vitrolles, France) with forepaws and then pulled backward by the base of its tails in the horizontal plane. At the grasp release, the force applied to the bar was recorded as the maximum strength of the grip. Each animal was tested 5 consecutive times with 1 min break between the tests. The grip strength was presented as the mean value of the 5 tests and weight-normalized grip strength (g force per g body weight).

### DEXA scan

After the tumor tissue was harvested, the animal carcasses were fixed in 10 % formalin for one week and then transferred to 70% ethanol for storage until DEXA scan was performed. On the day of the scan, fixed tissue was air-dried for 2 hours prior to DEXA procedure. The scanning and subsequent analysis, including fat percent, lean percent, and bone mineral content (BMC) percent, were carried out using InAlyzer2 (Micro Photonics Inc., Allentown, PA, US)

### Sample preparation and LC-MS Assay

Whole blood samples were collected through cardiac puncture upon the endpoint of the above-mentioned voluntary stretching/voluntary running experiment and centrifuged at 2000g for 10 min to collect plasma. Following centrifugation, separated plasma was then stored in a -30°C freezer until use. Proteomics analysis of plasma was conducted using a magnetic bead-based ENRICH-iST kit (PreOmics, Planegg, Germany) in accordance with the manufacturer’s instructions. Briefly, 20 µL of plasma sample was transferred to reaction tubes to incubate with magnetic beads on a thermoshaker (Themomixer C, Eppendorf, Enfield, CT, USA) at 30 °C and 1200 rpm for 30min, followed by 10-min lysis at 95 °C and 3-hr enzyme digestion at 37 °C then peptide cleanup in the cartridge at RT. Nano LC-MS/MS analysis of tryptic peptides was carried out with a Thermo Scientific Fusion Lumos tribrid mass spectrometer interfaced with an UltiMate3000 RSLCnano HPLC system. For each analysis, 1 µg of the tryptic digest was loaded and desalted in an Acclaim PepMap 100 trapping column (75 µm×2 cm) at 4 µL/min for 5 min. Peptides were then eluted into an Aurora Ultimate TS 25×75 C18 UHPLC column (Ionopticks, Australia) 75 µm×250 mm and chromatographically separated using a binary solvent system consisting of A: 0.1% formic acid and B: 0.1% formic acid and 80% acetonitrile at a flow rate of 300 nL/min. A gradient was run from 1% B to 32%B over 110 minutes, followed by a 5-minute wash step with 90% B and a 14-minute equilibration at 1% B before the next sample was injected. Precursor masses were detected in the Orbitrap at R=120,000 (m/z 200) in profile mode. Fragment masses were detected in the Orbitrap at R=15,000 (m/z 200) in centroid mode. Data-independent MSMS was carried out for m/z 400-1000 with 13 m/z windows. AGC target was set at 200%, and the maximum injection time was 20 ms. Protein identification and quantification were carried out by Spectronaut 19 using Direct DIA with software default settings. Raw data was searched against a mouse protein database from Uniprot (2022.10.20). Carbamidomethylating of C was set as a fixed modification, and oxidation of M was set as a variable modification. Quantification was based on the MS2 area.

## Supplementary figures

**Fig. S1.**
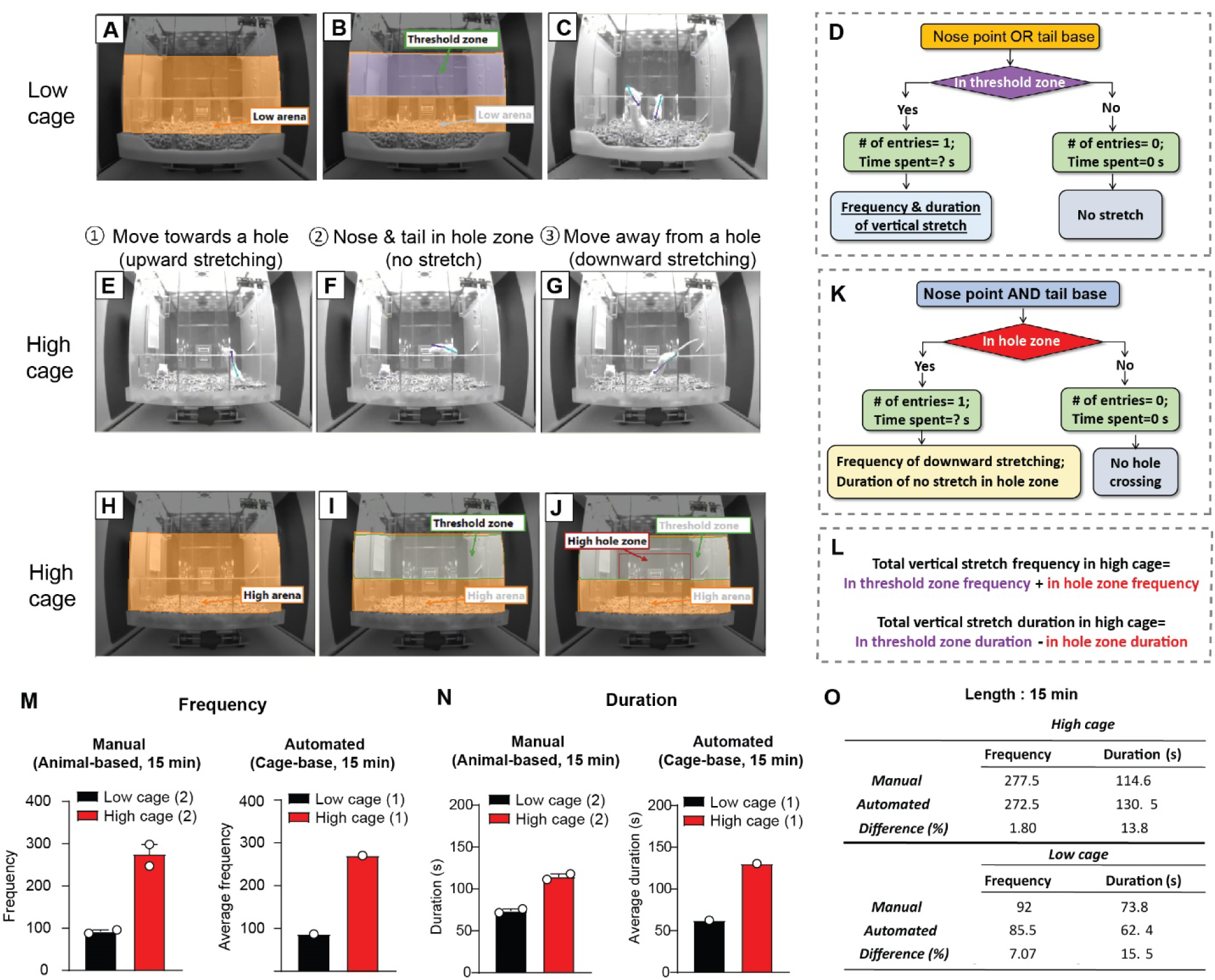
Vertical stretch monitoring and behavior analysis strategy. (**A, B**) The arena setting (A) and the specific threshold zone (B) defined to analyze vertical stretching behavior in the low cage. (**C**) An example image showing the nose point and tail-base point with a connecting line displayed during behavior tracking. (**D**) A flow chart of the vertical stretching tracking algorithm in the low cage. (**E-G**) Example images showing the movement breakdowns of the hole crossing behavior in the high cage. (**H-J**) In the arena of high cage (H), besides a threshold zone (I), a “high hole zone” (J) was specifically defined to analyze the vertical stretching occurs around the divider holes in the high cage. (**K**) A flow chart for the additional algorithm to analyze the total frequency and duration of vertical stretching in the high cage zone. **(L)** Equations for calculating the total frequency and duration of vertical stretching in high cage. (**M-N**) Comparison of results of manual analysis versus software-based automated analysis of the frequency (M) and duration (N) of the events of vertical stretching in low cage and high cage in a 15-minute video footage from the pilot study. Animals in the high cage stretched three times more frequently and nearly twice as long as those in the low cage (manual scoring: animal-based, n=2/group; automated scoring: cage-based, n=1/ group). (**O**) A summary of the difference between manual vs. automated analysis of frequency and duration in low and high cages. The tracking errors between the automated and manual scoring would not significantly impact the software’s ability to detect the difference in vertical stretching performance between the experimental and control groups.

**Fig. S2.**
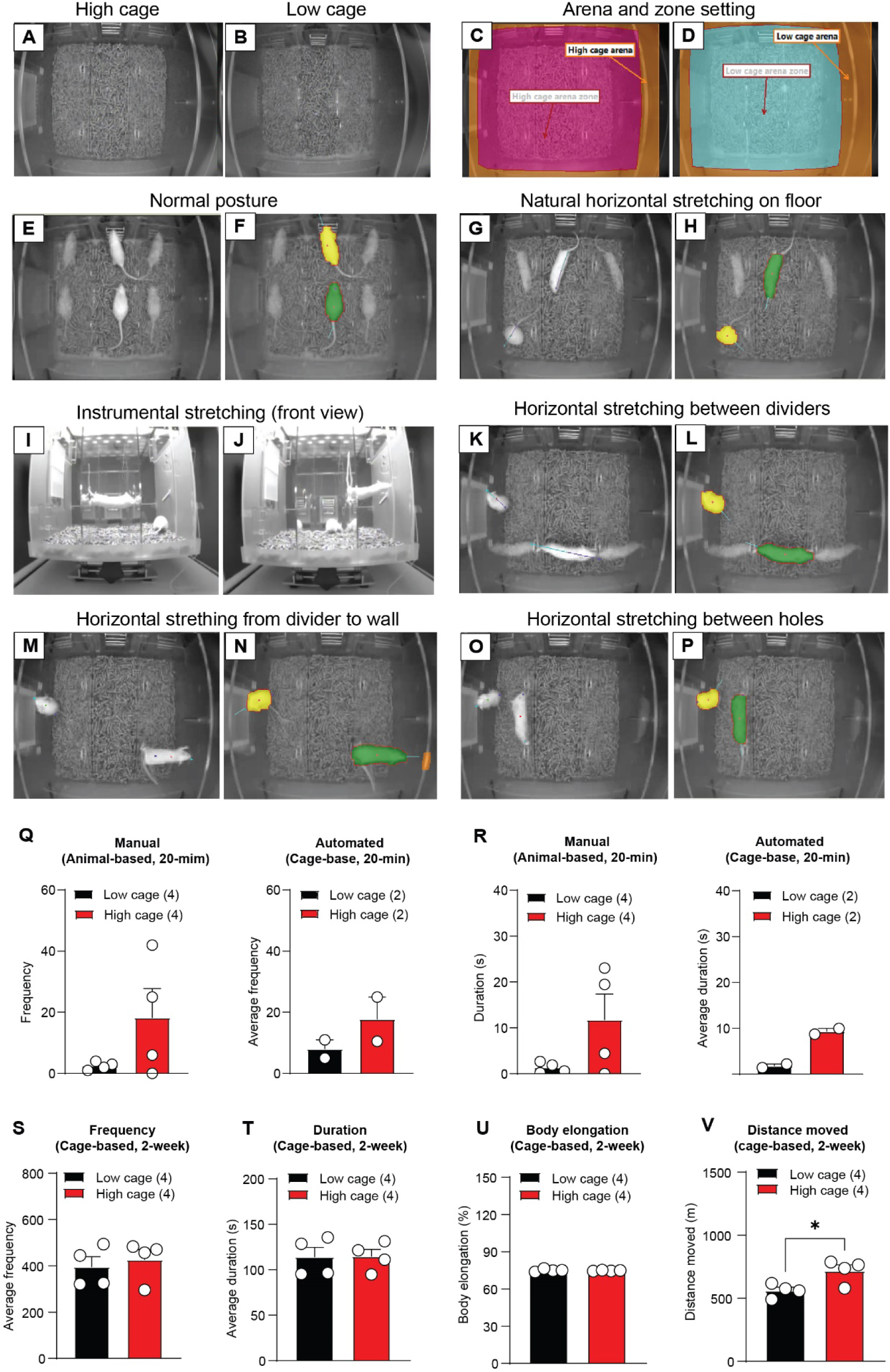
Analysis of horizontal stretching and locomotion. **(A-D)** Top-view images of high cage (A) and low cage (B), along with the arena and zone configuration in both cages in EthoVision XT 17 software (C, D). (**E-H**) Example images illustrating normal postures (no stretch, E, F) and natural horizontal stretching on the floor level (G, H) in two animals, with distinct color fills assigned by the software to distinguish their identities (F, H). (**I, J**) Front-view images showing instrumental horizontal stretching elicited by the elevated divider holes in the high cage, occurring between two cage dividers (I) as well as divider and wall (J). **(K-P)** Top-view images showing the different forms of detected horizontal stretching occurring between dividers (K, L), between a divider and wall (M, N) and between two holes on the same dividers (O, P). **(Q-R)** Comparison between the automated analysis and manual scoring of horizontal stretching for a 20-min video section in the pilot study showing that high cage setting also increases frequency (Q) and duration (R) of horizontal stretching (Q, frequency: manual scoring, animal-based, n=4 /group, low cage vs. high age, *p*=0.15; automated scoring, cage-base, n=2/group; R, duration: manual scoring, animal-based, n=4/group, *p*=0.12; automated scoring, cage-base, n=2/group) **(S-V)** A two-week long-term observation of horizontal stretching did not detected difference in horizontal stretching frequency (S), duration (T) or average body elongation (U, body elongation percent) between tumor-bearing animals in low cages and high cages, but an increase in locomotion in high cage animals (V) in breast tumor model experiment (S, 2-week frequency, cage-based, n=4/ group, *p*=0.64; T, 2-week duration, cage-based, n=4/group, *p*=0.95; U, body elongation percent, cage-based, n=4/group, *p*=0.69; V, distance-moved, cage-based, n=4/group, *p*=0.04). The data are expressed as means ± SEM. Statistics: (Q) and (R) manual scoring: *F*-test from nested mixed model; (S) to (T): Satterthwaite t-test. [\**p*<0.05].

