## Supplementary figures and images for "A novel mouse model of voluntary stretching and its application in breast cancer research"

### Fig. S1

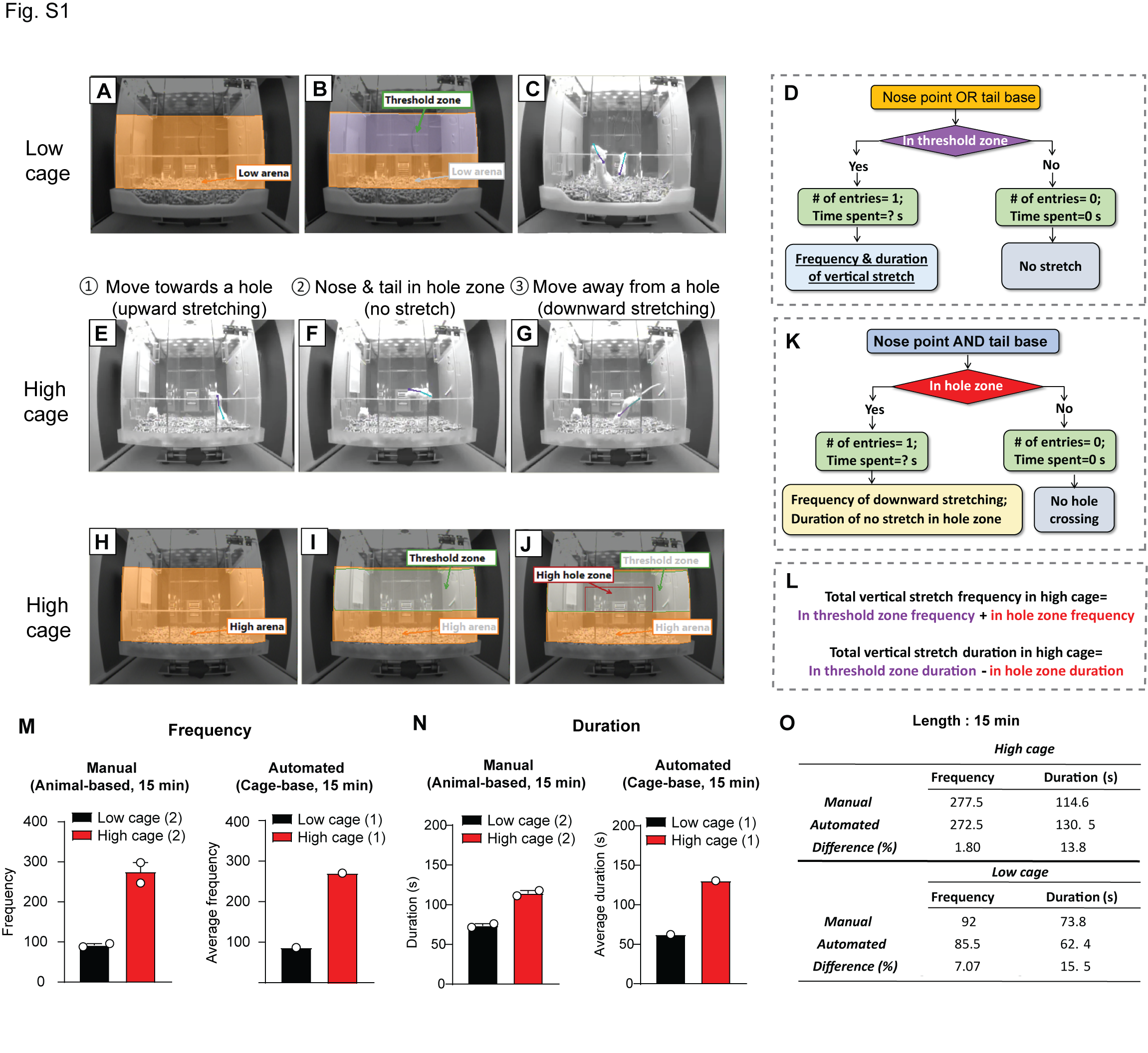

### Fig. S2

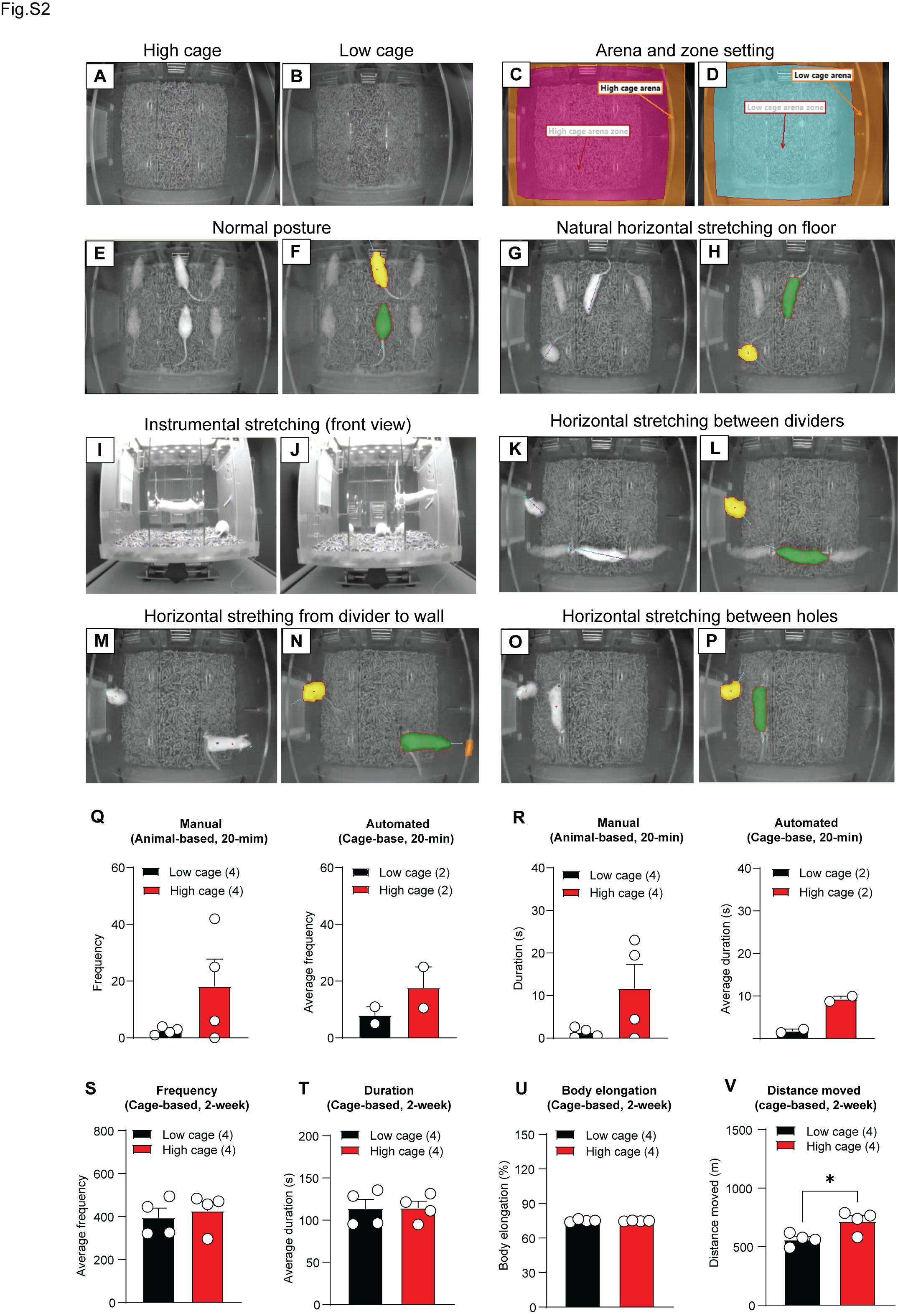
